# Biotext: Exploiting Biological-Text Format for Text Mining

**DOI:** 10.1101/2021.04.08.439078

**Authors:** Diogo de Jesus Soares Machado, Camilla Reginatto De Pierri, Letícia Graziela Costa Santos, Leonardo Scapin, Antonio Camilo da Silva Filho, Camila Pereira Perico, Fabio de Oliveira Pedrosa, Roberto Tadeu Raittz

## Abstract

The large amount of existing textual data justifies the development of new text mining tools. Bioinformatics tools can be brought to Text Mining, increasing the arsenal of resources. Here, we present BIOTEXT, a package of strategies for converting natural language text into biological-like information data, providing a general protocol with standardized functions, allowing to share, encode and decode textual data for amino acid and DNA. The package was used to encode the arbitrary information present in the headings of the biological sequences found in a BLAST survey. The protocol implemented in this study consists of 12 steps, which can be easily executed and/ or changed by the user, depending on the study area. BIOTEXT empowers users to perform text mining using bioinformatics tools. BIOTEXT is freely available at https://pypi.org/project/BIOTEXT/ (Python package) and https://sourceforge.net/projects/BIOTEXTtools/files/AMINOcode_GUI/ (Standalone tool).

## 1. INTRODUCTION

Texts are the most natural way of storing information and are responsible for the most extensive inventory of world scientific knowledge (TSHITOYAN et al., 2019). In sharp contrast to the analysis options, the mass of textual information in public databases demonstrates the need to develop tools to meet the growing demand for data manipulation and analysis (KOCH et al., 2020). For the omics area, whose primary focus is the analysis of complex biological events, Text Mining (TM) combined with bioinformatics techniques represents an advance for exploring and analyzing unstructured data (ZAFEIROPOULOS et al., 2022).

Today’s TM tools address different methodologies, with potential for research in the most diverse scenarios (TABLE S1 – for details of review, see FILE S1). These approaches involve the main tasks such as Information Retrieval (IR), Named Entity Recognition (NER), and Information extraction (IE) (FIGURE S1), and a series of algorithms can be run depending on the type of analysis (TABLE S2) (FLEUREN; ALKEMA, 2015). However, we note that even with several aspects of TM well resolved in the literature, it is still possible to implement these processes advantageously. One of these ways is directly transposing Bioinformatics technologies to the TM area.

The exploration of biological sequences has a strong analogy with TM, since it works fundamentally with character string analysis (HARMSTON et al., 2010). From more traditional techniques such BLAST for sequence alignment (ALTSCHUL et al., 1990), to more recent ones such as the representation of biological sequences in vector spaces (ASGARI, MOFRAD, 2015; LEIMEISTER et al., 2018; DE PIERRI et al., 2020; RAITTZ et al., 2021) can be brought to the TM area, adding to the arsenal of existing text analysis tools. In addition, as in the field of Bioinformatics, vector representation of texts is also a trend in text processing (MIKOLOV et al., 2013; LEVY, GOLDENBERG, 2014; PENNINGTON et al., 2014), since TM vectoral technologies are equally promising (MA et al., 2015; LILLEBERG et al., 2015; DEVLIN et al., 2018; HASSANI et al., 2020).

However, the Bioinformatics tools available today do not process normal texts. It is necessary to use the Biological Sequence Format (BSF) to perform analysis. Thus, here we present an approach that adapts texts in natural language to be processed with several Bioinformatics tools, including the main resources for the comparison of sequences added to the agility and robustness of vectorization technologies. BIOTEXT is a package to perform text mining strategies using bioinformatics tools, which provides AMINOcode and DNAbits features to encode text in BSF.

We explored tools for encoding arbitrary information present in the headers of biological sequences found in a typical BLAST search. We demonstrate that texts can be encoded into biological information using computational strategies based on bioinformatics. The benefit of representing texts in the form of biological sequences can be extended to the level of physical information storage, as the use of DNA for this task allows the storage of information in high density in much smaller volumes (TSHITOYAN et al., 2019; KOCH et al., 2020).

## 2. METHODS

### 2.1 BIOTEXT Implementation

The Python package is available from the PyPI repository at https://pypi.org/project/BIOTEXT/. The case study is accessible through the script “fastaHeader2plot.py, available at https://github.com/diogomachado-bioinfo/BIOTEXT-examaples/tree/master/fastaHeader2plot. In addition, a complete list containing all BIOTEXT *functions is available in Table S3*.

### 2.2 BIOTEXT application

To demonstrate one of the applications of BIOTEXT using traditional methods in bioinformatics, we explored the local alignment for direct comparison between texts. We randomly selected the protein hypothetical “WP_011156533.1” of a set of sequences from a previous study (RUBEL et al., 2015) for BLAST research, using default parameters; the first 100 hits were considered (for details, see TABLE S4).

## 3. RESULTS AND DISCUSSION

### 3.1 Converting texts into biological sequence format (BSF)

The BIOTEXT package includes two functions – and respective reverse functions - to code conventional texts to a valid biological format.

AMINOcode consists of replacing text characters with letters used to represent amino acids, according to a coded list. Table 1 shows the character substitution rules in AMINOcode. In addition, we present a minimal and an expanded encoding table that includes number and punctuation details representations. Whereas the 20 letters used to represent the amino acids are less than 26 needed to represent the whole alphabet, we define a conversion rule. For consonants, except for W and Y, the replacement is carried by a single letter of the amino acid representation (TABLE1). The coding replaces the vowels with two characters. The first is Y, a ‘key’ that defines the subsequent letter. Next, in the same perspective, all kinds of spaces in the texts are coded with YS. Finally, we propose the expanded version for digits and punctuation. In the expanded one, we use a key with two key characters, YP followed by the coding type. The minimal version can cause slight information, but it may be helpful in some instances. Also, we define a wildcard representation, YK, for any exception to Table 1.

**TABLE 1.**
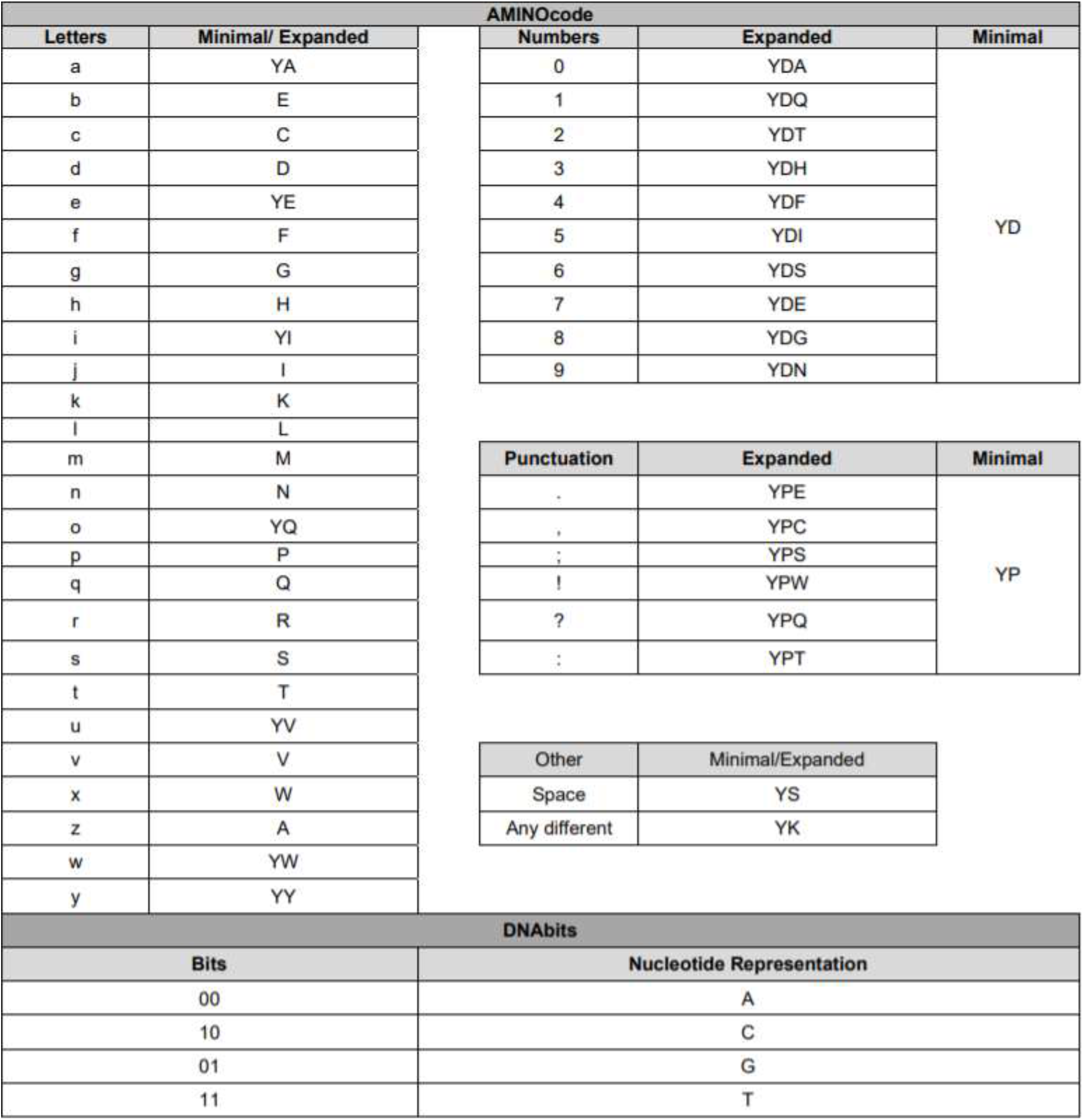
AMINOcode and DNAbits Dictionary. Legend: DNAbits dictionary is represented by the nucleotides A (adenine), C (cytosine) G (guanine) and T (thymine);

DNAbits performs the conversion of text characters splitting every byte into four pairs of bits. The codding maintains the information according to the “American Standard Code for Information Interchange” (ASCII), replacing every couple of bits by A, C, G, and T according to the predefined rule (Table 1). In DNAbits, the characters are encoded from binary form by the function in Eq.(1), and the decoding of the DNAbits is translatable to the original text.

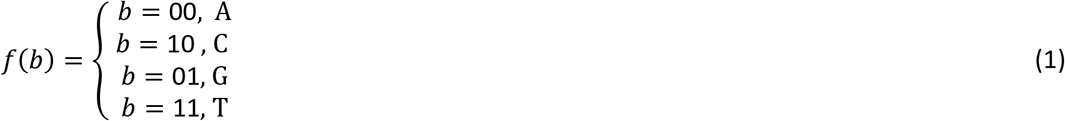

where *b* is any pair of bits, for example, the binary form for the character representing the letter ‘a’ is 10000110, resulting in CAGC after DNAbits encoding.

### 3.2 Interpreting BLAST results

We randomly selected the hypothetical protein WP_011156533.1” from a set of sequences from a previous study (RUBEL *et al*., 2015) for research on BLAST (ALTSCHUL *et al*., 1990) and run with the default parameters; the first 100 hits were considered (TABLE S4). Next, the headers of the sequences were converted to BSF using AMINOcode, resulting in a FASTA file containing the text information. For the header’s contents analyses, the headers sequences were converted to vectors by the SWeeP function (DE PIERRI *et al*., 2020) Then, the headers groups were defined by agglomerative clustering (FIGURE 1a). Finally, we aligned the sequences from each cluster separately with Clustal Omega (SIVIERS *et al*., 2018) and obtained the sequence consensus decoded through reverse AMINOCode, as shown in Figure 1b. The entire process consists of 12 steps (see Figure S2):

1. Extract headers from files;
2. Use expanded version with number and punctuation details (optional);
3. Find and remove patterns using regular expression for ID from all encoded texts;
4. Vectorize encoded header in a matrix with a line for each sequence;
5. Obtain the first two dimensions of loading of the PCA from SWeeP matrix;
6. Hierarquical clustering for two dimensions of loading, using the agglomerative clustering (scikit-learn Python library);
7. Agglomerative clustering, using the 0,015 threshould (optional);
8. Group the encoded texts according to the clusters;
9. Obtain the consensus of sequences in each cluster from Clustal Omega alignment;
10. Find and remove patterns using regular expression for species name from consensus strings;
11. Decode the profile strings in texts;
12. Plot a 2D graph with two loading dimensions with the equivalent texts derived from the profiles as a legend.

**FIGURE 1.**
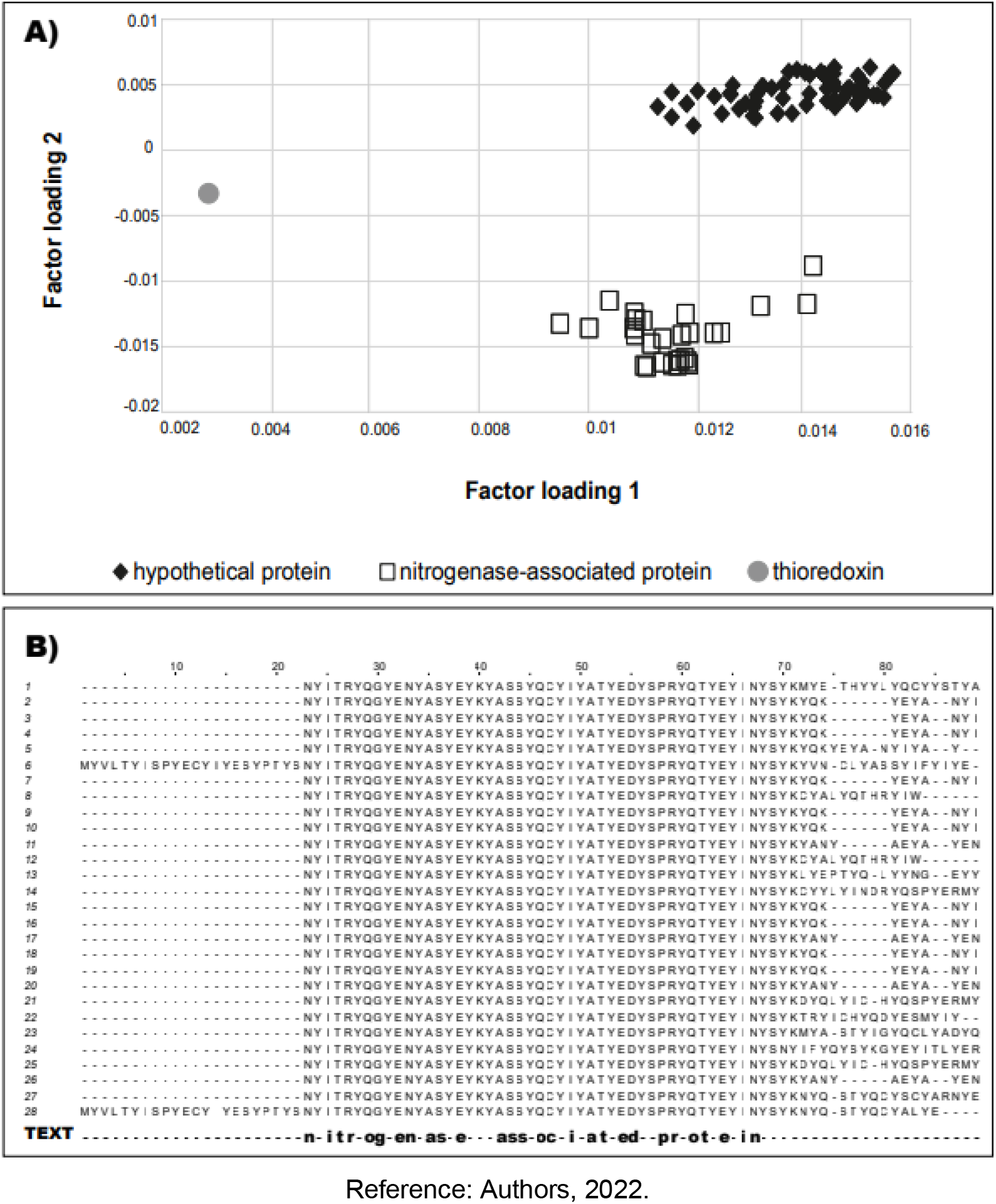
Graphic representation of sequence headers. a) Each dot represents a header obtained by the BLAST search using the WP_011156533.1 protein as a query. Consensus Headers are defined by: Hypothetical protein (lozangue), Nitrogenase-associated protein (square) and Thioredoxin (circle). The PCA was generated using the scikit-learn library (PEDREGOSA *et al*., 2011). b) Representations of alignments with coded texts from a cluster. Alignment positions are shown at the top of the figure. Cuts were made in the alignments to highlight the region of greatest importance. Headers are displayed with identification numbers for each sequence (left). The last line, represented by the “TEXT” header is the decoded consensus text, properly aligned according to the coded texts.

According to the exploited data, the use of expanded or minimal versions is optional. Steps 3 and 10 correspond to removing unique patterns from each header (ID and species), which do not provide helpful information for the clustering and analysis of the texts.

Figure 1a shows the graph of the two main components of a vector PCA that represent the header sequences. Clusters are labeled by the consensus header automatically generated for the clusters: “Hypothetical protein”, “Nitrogenase-associated protein,” and “Thioredoxin”. Distances among points (graph in FIGURE 1a) result from variations in the headers’ text. The more variations between texts, the further apart the dots appear on the chart. Even points of the same cluster are not in the same position, as the texts represented are not identical. The exploitation of the BSF provided by BIOTEXT functions allowed us to derive information from a hypothetical BLAST query automatically. Interpreting a hypothetical protein is a challenge in bioinformatics, and comparisons with various known sequences are often adopted to understand them (IJAQ *et al*., 2019). We seek to explore the relationship of information present in the header of a hypothetical protein using standard bioinformatics tools to accomplish this task.

### 3.3 BIOTEXT application experiment in large corpus

With the string “thioredoxin”, we searched on PubMed using the Biopython package “Bio.Entrez” (COCK *et al*., 2009) (on March 18, 2022), resulting in 13,734 entries. We extract only the titles retrieved from the search and apply AMINOcode encoding using the expanded version (with detailed digits and punctuation). Then, the coded texts were vectorized and derived in a dendrogram with Euclidean distance based on the distance matrix - implementation of the SciPy library (VIRTANEN *et al*., 2020). We use the scikit-bio library to transform the dendrogram into Newick format. The resulting Newick file, input text, and script to re-run all processes are available at https://github.com/diogomachado-bioinfo/BIOTEXT-examples/tree/master/titleSet2tree.

It is essential to highlight that the execution was quickly on an ordinary computer (i5-3470 processor and 16 Gb of DDR3 RAM). The entire process took 317.64 seconds. In particular, encoding with AMINOcode took 1.12 seconds, vectoring with SWeeP 244.38 seconds and creating the dendrogram 72.14 seconds.

## 4. CONCLUSION

Here we present BIOTEXT and run an example in interpreting a result of BLAST. The library can be applied in various issues involving word processing and text mining, employing FASTA files and Bioinformatics tools. In addition, this study can contribute to research on the vector representation of texts.

We encode texts (FASTA headers) in BSF and, therefore, could be treated similarly to what would be with biological FASTA. The case study shows that it is easy to perform TM tasks directly in BSF coded texts using bioinformatics tools and strategies, such as Clustering, PCA, alignment, and BLAST for consensus identification and even pattern recognition or machine learning. However, the fact that the available tools are dependent on vocabulary makes text handling complex. BIOTEXT incorporates compact vector technology to streamline analysis and eliminate the need for predefined vocabulary.

BIOTEXT combined with SWeeP vectors technology provided handling of large amounts of text, with fast execution time even on a personal computer. Moreover, our proposal is versatile, allowing the use of the entire arsenal of algorithms/methods in bioinformatics for analyzes in which texts are transformed into BSF. BIOTEXT is a general purpose tool for text mining that allows one to use different bioinformatics algorithms for multipurpose analyses and perform the main text mining tasks (IR, NER, and IE). In short, our approach can be handy for bioinformatics, given the analysts’ experience with these features.

## ACKNOWLEDGMENT

The authors thank the group of Artificial Intelligence Applied to Bioinformatics of the Federal University of Paraná and the Coordination of Improvement of Higher Education Personnel – CAPES for the support.

