## Supplementary Information for "Biotext: Exploiting Biological-Text Format for Text Mining"

TABLE S1.Text mining tools and approaches for Omics sciences

| Tool | Description | Purpose | Task | Input | Input | Authors and Year |
|---|---|---|---|---|---|---|
| MedMiner | Web-based tool that filters the literature and returns organized information. | Biocuration | IR | List of genes | List of genes | TANABE *et al.*, 1999 |
| STRING | Tool for recurring instances of neighboring genes, retrieving and displaying the genes that in clusters on the genome. | Biocuration | IR | Query | Query | SNEL *et al.*, 2000 |
| MeshMap | Text mining system to exploit the MeSH indexing accompanying the Medline records and compare entities of the same type. | General purpose | IR NER | Query | Query | SRINIVASAN, 2001 |
| Pubmatrix | Comparison tool for systematic and automatic search on PubMed. | General purpose | IR | Query | Query | BECKER *et al*, 2003 |
| Chillibot | NLP-based text-mining approach which construct content-rich relationship networks among biological concepts, genes, proteins, or drugs. | General purpose | IR NER IE | List of genes | List of genes | CHEN; SHARP, 2004 |
| Textpresso | A text Mining tool that splits papers into sentences, and sentences intowords or phrases, including all categories of the Gene Ontology database | General purpose | IR NER IE | Texts | Texts | MULLER *et al.*, 2004 |
| BioCreAtIvE | A multitasking Text Mining tool to assess the state of the art applied tobiological problems. | General purpose | IR NER IE | Texts | Texts | HIRSHMAN *et al.*,2005 |
| TAXONGRAB | A tool for extract taxonomic names from English, Spanish, French and German texts, using a lexicon of words. | Taxonomy | NER | Texts | List of scientific names | KONING *et al.*, 2005 |
| CRF | A tool for tagging gene and protein mentions from text using the probabilistic sequence tagging framework. | General purpose | NER | List of terms | List of terms | MACDONALD; PEREIRA, 2005 |
| ABNER | A tool for molecular biology text mining based on machine learning system to extract features and tag entities. | General purpose | NER | List of terms | List of terms | SETTLES, 2005 |

(Continue)

| | | | | | | |
|---|---|---|---|---|---|---|
| AliBaba | An interactive tool for graphical summarization parsing Abstracts of PubMed searches. | General purpose | IR | Query | Query | PLAKE *et al.*, 2006 |
| STITCH | A TM tool for interactions of chemicals that integrates information about interactions from metabolic pathways, crystal structures, binding experiments and drug–target relationships. | Chemicals | IR<br>NER<br>IE | Query | Query | KUHN *et al.*, 2007 |
| Whatizit | A Tool for tag concepts in texts based on thesaurus. | General purpose | NER | Texts | Texts | REBHOLZ-SCHUHMANN *et al.*,2007 |
| TaxonFinder | A tool designed to find scientific names from text with the help of separate dictionaries for species and genus names | Taxonomy | NER | Texts | List of scientific names | LEARY *et al.*, 2007 |
| PolySearch | Web-tool system which makes relationships between several queries in the biomedical sciences. | General purpose | IR<br>NER<br>IE | Query | Query | CHENG *et al.*, 2008 |
| E3Miner | Tool that extracts comprehensive knowledge about the ubiquitin-protein ligase (E3) from the scientific literature. | Molecular function | IR<br>IE | Abstracts | Abstracts | LEE *et al.*, 2008 |
| ANNE O'TATE | An integrated, generic tool for summarization, drill-down and browsing of PubMed search results | General purpose | IR<br>NER | Query | Query | SMALHEISER *et al.*, 2008 |
| MedlineRanker | Allows flexible and fast ranking of Medline abstracts for a topic of interest without expert knowledge. | General purpose | IR | Query | Query | FONTAINE *et al.*, 2009 |
| MiSearch | Search tool that ranks citations based on a statistical model for the likelihood that a user will choose to view them. | General Purpose | IR | Query | Query | STATES *et al.*, 2009 |

(Continue)

| | | | | | | |
|---|---|---|---|---|---|---|
| GoGene | Performs high-throughput text mining to complement annotation of genes. | Biocuration | IR<br>IE | List of Genes | List of Genes | PLAKE *et al.*, 2009 |
| @Note | Its main functional contributions are the ability to process abstracts and full-texts. | General Purpose | IR | Abstracts / full-texts | Abstracts / full-texts | LOURENÇO et al., 2009 |
| PLAN2L | A web-based tool that integrates text mining and information extraction techniques to access useful information for analyzing genetic, cellular andmolecular aspects of Arabidposis thaliana. | Molecular function | IR<br>NER<br>IE | Query | Query | KRALLINGER *et al.*,2009 |
| PathBinder | Plant-based tool to explore literature, and to annotate metabolic andregulatory interactions and biomolecules. | Molecular function | IR<br>NER<br>IE | Query | Query | ZHANG *et al.*, 2009 |
| RefMed | A multi-level relevance feedback system for PubMed. | General Purpose | IR | Query | Query | YU *et al.*, 2010 |
| LINNAEUS | A system that uses a dictionary and a set of regular expressions to identify species mentions in text. LINNAEUS uses dictionaries for scientific and common names to construct a DFA (Deterministic Finite Automaton). | Taxonomy | NER | Texts | Query<br>List of scientific names | GERNER *et al.*, 2010 |
| LAITOR | A toll for analyze co-occurrences of bioentities, biointeractions, and other biological terms in MEDLINE abstracts. | Molecular function | IR<br>NER | Query | Query | SILVA *et al.* 2010 |
| DataShield | Allows the analysis of sensitive individual-level data from one study. It can be applied to post-publication sensitive data analysis or text data analysis. | General Purpose | IR<br>NER | Texts | Texts | WOLFSON *et al.*, 2010 |
| BSQA | Web server that performs integrated text mining, recognizing several entities and relations in Medline documents about Drosophila melanogaster. | Molecular function | IR<br>NER | Query | Query | HE *et al.*, 2010 |

(Continue)

| | | | | | | |
|---|---|---|---|---|---|---|
| EMU | A Text mining tool for identify mutations and their associated genes. | Molecular function | IR<br>NER<br>IE | Texts | Texts | DOUGHTY *et al.*,2010 |
| Quertle | Allows a semantic search in multiple biomedical databases and runs aquery via relationships between concepts. | General Purpose | IR<br>NER<br>IE | Query | Query | GIGLIA, 2011 |
| GeneValorization | A web-based tool that gives an overview of the bibliography available | Molecular function | IR<br>NER | List of Genes | List of Genes | BRANCOTTE *et al.*, 2011 |
| FACTA+ | Real-time text-mining system for finding and visualizing indirect associations between biomedical concepts from MEDLINE abstracts. | General Purpose | IR<br>NER<br>IE | Query | Query | TSUROUKA *et al.*,2011 |
| OrganismTagger | A hybrid rule-based/machine learning system to extract organism mentions from the literature | Taxonomy | NER | List of names | Query<br>List of scientific names | NADERI *et al.*, 2011 |
| BeeSpace Navigator | Tool for exploratory analysis and extract gene functions. | Molecular function | IR<br>NER<br>IE | Query | Query | SARM, *et al.*, 2011 |
| PESCADOR | A web tool that extracts a network of interactions from a set of PubMedabstracts, filtering according to user-defined concepts. | General purpose | IR<br>NER<br>IE | Query | Query | SILVA *et al* 2011 |
| OSCAR4 | The tool recognizes sections of reported experimental and use regular expressions to match the journal-mandated formats in which are reportedin the literature. | Chemicals | IR<br>NER | text | text | JESSOP *et al.*, 2011 |
| BioContext | TM system which extracts, extends and integrates results from a number of tools performing entity recognition, biomolecular event extraction and contextualization. | Molecular function | IR<br>NER | Query | Query | GERNER *et al.* 2012 |

(Continue)

| | | | | | | |
|---|---|---|---|---|---|---|
| NetiNeti | A tool focused on recognition/discovery of scientific names of organisms from various text sources. | Taxonomy | NER | Texts | List of scientific names | AKELLA *et al.*, 2012 |
| MyMinner | MyMiner is a free and user-friendly text annotation tool aimed to assist in carrying out the main biocu ration tasks and to provide labelled data for the development of text mining systems. | Biocuration | NER | Texts | Texts | SALGADO *et al.* 2012 |
| BioC | BioC allows a large number of different annotations to be represented such as genes or diseases and relationships between named entities. | General Purpose | IR<br>NER | Text | Text | COMEAU *et al.*, 2013 |
| PubTator | Web server tool for text mining that allows multiple challenge-winning to assist biocuration, by featuring a PubMed-like interface. | Biocuration | IR<br>NER | Query | Query | WEI *et al.*, 2013 |
| SPECIES | Dictionary-based approaches, being able to recognize vernacular names in text in addition to scientific ones. | Taxonomy | NER | Texts | List of scientific names | PAFILIS *et al.*, 2013 |
| EVEX | Is a text mining resource which focuses on biomedical event extraction and gene interactions, covering the whole biomedical literature available in PubMed. | Molecular function | IR<br>NER<br>IE | Query | Query | LANDEGHEM *et al.*,2013 |
| BeCAS | It's a web-based tool for annotation of biomedical concepts that can be integrated on larger text-processing pipelines or incorporated on externalweb pages. | General Purpose | NER | Abstracts | Abstracts | NUNES *et al.*, 2013 |
| OntoGene | A TM system specialized in the detection of named entities as proteins,genes, and others, allowing a deeper connection between the unstructured information. | Molecular function | IR<br>NER<br>IE | Query | Query | RINALDI *et al.* 2014 |
| Taxamatch | Employs a custom distance algorithm in tandem with a phonetic algorithm,together with a rule-based approach to produce improved levels of recall, precision and execution time over the n-grams algorithms. | Taxonomy | IR<br>NER | Query | Query<br>List of scientific names | REES, 2014 |

(Continue)

| | | | | | | |
|---|---|---|---|---|---|---|
| Alkemio | A tool that ranks all annotated chemicals (including drugs) in the literature for any biomedical topics. | Chemicals | IR NER | Query | Query | GIJÓN-CORREAS et al., 2014 |
| PubstractHelper | Allows the researcher to put up to ten different keywords and highlight sentences with co-occurring keyword in different colors. | General purpose | IR | Query | Query | CHEN; HO, 2014 |
| RLIMS-P 2.0 | Rule-based text-mining program specifically designed to extract protein phosphorylation information on protein kinase,substrate andphosphorylation sites from biomedical literature. | Molecular function | IR NER IE | Texts | Texts | TORII *et al.*, 2015 |
| miRTex | Text-mining system that extracts miRNA-gene regulation relations andmiRNA-target relations from literature | Molecular function | IR NER IE | Texts | Texts | LI *et al.*, 2015 |
| OntoMate | Ontology-driven system used as a replacement for the PubMed search engine in the Rat Genome Database that works on the gene curation workflow. | Molecular function | IR NER IE | Abstracts | Abstracts | LIU *et al.* 2015 |
| DISEASES | A system for extracting disease–gene associations from biomedical abstracts, using a dictionary-based tagger for named entity recognition ofhuman genes and diseases. | Molecular function | IR NER IE | Abstracts | Abstracts | PLETSCHER-FRANKILD *et al.*,2015. |
| Aggregator | Machine learning approach to identify articles that derived from the sameclinical trial. | General Purpose | IR NER IE | Train sets | Train sets | SHAO *et al.*, 2015 |
| NOBLE | A recognition component for NLP (Natural Language Processing) pipelines. | General Purpose | NER | Query | Query | TSEYTLIN *et al.*, 2016 |
| @Minter | An automated information extraction system based on Support Vector Machines to analyze paper abstracts and infer microbial interactions. | Molecular function | IR NER IE | Abstracts | Abstracts | LIM *et al.*, 2016. |

(Continue)

| | | | | | | |
|---|---|---|---|---|---|---|
| MiRiaD | A text mining tool that automatically extracts associations between microRNAs and diseases from the literature. | Molecular function | IR NER IE | Query | Query | GUPTA *et al.*, 2016 |
| SparkText | Text mining framework, on a Big Data infrastructure, which is composed of Apache Spark data streaming and machine learning methods. | General Purpose | IR NER IE | Abstracts/full-texts | Abstracts/full-texts | YE *et al.* 2016 |
| LimTox | An online text mining application that automatically extracts toxicology relevant information from text, with special emphasis on drug-inducedadverse hepatobiliary reactions. | Chemicals | NER | Preprocessed documents | Preprocessed documents | CAÑADA *et al.*, 2017 |
| PaperBlast | PaperBLAST uses Europe PMC to search the full text of scientific articles for references to genes, to find similar proteins presents in snippets of text. | Biocuration | IR | Query | Query | PRICE *et al.*, 2017. |
| EGenPub | Selects articles about a certain protein and identify an additional bibliography from the UniProt knowledgebase, based in SVM. | Molecular function | IR NER IE | UNIPROT accession number | UNIPROT accession number | DING *et al*, .2017. |
| BELMiner | Semi-automated Text minig system used to facilitate the curation tasks on building biological networks from unstructured scientific texts. | General Purpose | IR NER IE | Query | Query | RAVILKUMAR *et al.*,2017 |
| Nala | Nala capture natural language and mention of the target pattern mutation by combining conditional random fields with word incorporation featureslearned without supervision. | Molecular function | IR NER IE | Texts | Texts | CAJUELA *et al.*, 2017 |
| Egard | A natural language processing-based text mining to extracting genomicanomalies association with Response to Drugs. | Chemicals | IR NER IE | Abstracts | Abstracts | MAHMOOD *et al.*,2017 |
| EzTag | Allows users to perform annotation. EzTag provides training data interactively. It supports all PubMed abstracts and PMC open access articles. | General Purpose | IR NER | Texts | Texts | KWON *et al.*, 2018. |

(Continue)

| | | | | | | |
|---|---|---|---|---|---|---|
| ITextMine | A system which uses an automated workflow that integrates multiple text mining tools and run it on large-scale text for knowledge extraction. | General Purpose | IR NER IE | Text | Text | REN *et al.*, 2018 |
| LitVar | A tool for the accurate search of variants and related information fromunstructured human-related biomedical literature | Molecular function | IR NER IE | Query | Query | ALLOT *et al.*, 2018 |
| Adjutant | R-based tool to support topic discovery for systematic and literature reviews. | General Purpose | IR NER IE | Query | Query | CRISAN *et al.*, 2019 |
| AlvisIR | Text mining tool for extracting information about microbial biodiversity infood. | General Purpose | IR NER IE | Abstracts | Abstracts | CHAIX *et al.*, 2019 |
| PPIcurator | Tool for extracting comprehensive protein-protein interaction information. | Biocuration | IR NER IE | Query | Query | LI *et al.*, 2019 |
| BioReader | Enables users to perform classification of scientific literature by text mining-based classification of article abstracts | General purpose | IR NER IE | Abstracts | Abstracts | SIMON *et al.*, 2019 |
| OGER++ | Performs document annotation by parsing, entity recognition and serialization, supporting concept recognition. | General purpose | IR NER | Abstracts | Abstracts | FURRER *et al.*, 2019 |
| CoCoScore | An unsupervised counting approach with a machine learning approach based on distant supervision to score sentence-level co-mentions. | General purpose | IR NER IE | Abstracts | Abstracts | JUNGE; JENSES,2020 |
| RWRMTN | A TM tool to predict novel disease-associated miRNAs. | Molecular function | IR NER IE | Query | Query | LI; TRAN, 2020 |

(Continue)

| | | | | | | |
|---|---|---|---|---|---|---|
| OmixLitMiner | The tool automates the retrieval of literature from PubMed based on UniProt protein identifiers, gene names and their synonyms, combined with user defined contextual keyword search | General purpose | IR<br>NER<br>IE | UNIPROT accession number | Query | STEFFEN *et al.*, 2020 |
| Melodi Presto | A tool that links risk factors and diseases by identifying overlapping<br>elements between two query lists using semantic triples. | General purpose | IR<br>NER | Query | Query | ELSWORTH; GAUNT, 2020 |
| LAITOR4HPC | Uses high performance computing aiming to access an unlimited number of abstracts to create new interaction networks or improve the ones that already exist. | Molecular function | IR<br>NER<br>IE | Query | Query | PIERECK *et al.* 2020 |
| ThermoScan | A text mining approach for the identification of relevant thermodynamic data on protein stability from full-text articles | Molecular function | IR<br>NER<br>IE | Full-texts | Full-texts | TURINA *et al.*, 2021 |
| PREGO | PREGO combines text mining and data integration techniques to explore associations from data and metadata scattered in the scientific literature and in public omics repositories. | Taxonomy | IR<br>NER<br>IE | Taxon Query | Tab of Organisms | ZAFEIROPOULOS *et al.*, 2022 |

Reference: Authors, 2022.

*TABLE S2. Text mining algorithms/models and its purpose*

| Process | Type of Algorithms | Purpose |
|---|---|---|
| Information Retrieval | Tokenization | Break text into discrete words, called textual entities or tokens |
| | Stopping | Remove common words, such as “the”, “or” |
| | Steeeming | Remove prefixes and suffixes to normalize words |
| | Normalize case | Convert the text to either all lower or all upper case |
| Named Entity Recognition | Indexing | Processing a text to extract statistics considered important for representing the information and/or to allow fast search on its content. |
| | Dictionary-based methods | Matching a dictionary of terms, containing terms and synonyms, against the query text. |
| | TF-IDF | TF- IDF (Term Frequency - Inverse Document Frequency). stores text as weighted vectors; The frequency for a term is the number of times the term appears in a document. |
| | NLP methods | Low-level language processing and comprehension task; grammatical class tagging. |
| Information Extraction | Prediction models | Classifying of textual entities into predefined categories |
| | Clustering models | Clustering of textual entities into formerly undefined and unknown groups |
| | Association rules | Finding interesting association or correlation relationships among a large set of data items |

Reference: Authors, 2022.

TABLE S3. List of BIOTEXT functions

| Function Name | Description | Input | Output |
|---|---|---|---|
| BIOTEXT.aminocode.encodeText<br>BIOTEXT.aminocode.encodetext<br>BIOTEXT.aminocode.et | Encodes a string with AMINOcode. | text: natural language text string to be encoded;<br>detailing: details in coding. 'd' for details in digits. 'p' for details on the punctuation. 'dp' or 'pd' for both. | enc_text: encode text in string format. |
| BIOTEXT.aminocode.decodeText<br>BIOTEXT.aminocode.decodetext<br>BIOTEXT.aminocode.dt | Decodes a string with reverse AMINOcode. | text: text string encoded using the encodefile function to be decode;<br>detailing: details used in the text to be decoded. 'd' for details in digits. 'p' for details on the punctuation. 'dp' or 'pd' for both. | dec_text: decode text in string format. |
| BIOTEXT.aminocode.encodeFile<br>BIOTEXT.aminocode.encodefile<br>BIOTEXT.aminocode.ef | Encodes a text file or a list of strings with AMINOcode. | input_file_name: text file name or list of string. It can also be used in a list of SeqRecord*, in which case the function will automatically extract the headers to do the encoding; output_file_name: the name for the output file. If not defined, the result will only be returned as a variable;<br>detailing: same as in the encodetext function;<br>header_format: format for the headers of the generated FASTA. It can be 'number+originaltext', 'number' or 'originaltext'. 'number' is a count of the lines in the input file. Blank lines are considered in the count, but are not added to the FASTA file. 'originaltext' is the input text itself; verbose: if True displays progress. | records: list of SeqRecord;<br>if defined output_file_name a file will be saved. |
| BIOTEXT.aminocode.decodeFile<br>BIOTEXT.aminocode.decodefile<br>BIOTEXT.aminocode.df | Decodes a fasta file or a SeqRecord list with the reverse amino acid. | input_file_name: file name or list of SeqRecord;<br>output_file_name: the name for the output file. If not defined, the result will only be returned as a variable;<br>detailing: same as in the decodetext function;<br>verbose: if True displays progress;<br>output: string list. If defined output_file_name a file will be saved. | dec_list: string list;<br>if defined output_file_name a file will be saved. |
| BIOTEXT. dnabits.encodeText<br>BIOTEXT.dnabits.encodetext<br>BIOTEXT. dnabits.et | Encodes a string with DNAbits. | text: natural language text string to be encoded. | enc_text: encode text in string format. |
| BIOTEXT.dnabits.decodeText<br>BIOTEXT.dnabits.decodetext<br>BIOTEXT.dnabits.dt | Decodes a string with reverse DNAbits. | text: text string encoded using the encodefile function to be decode. | dec_text: decode text in string format. |

(Continue)

| | | | |
|---|---|---|---|
| BIOTEXT.dnabits.encodeFile<br>BIOTEXT.dnabits.encodefile<br>BIOTEXT.dnabits.ef | Encodes a text file or a list of strings with DNAbits. | input_file_name: text file name or list of string. It can also be used in a list of SeqRecord, in which case the function will automatically extract the headers to do the encoding; output_file_name: the name for the output file. If not defined, the result will only be returned as a variable;<br>header_format: format for the headers of the generated FASTA. It can be 'number+originaltext', 'number' or 'originaltext'. 'number' is a count of the lines in the input file. Blank lines are considered in the count, but are not added to the FASTA file. 'originaltext' is the input text itself;<br>verbose: if True displays progress. | records: list of SeqRecord.<br>if defined output_file_name a file will be saved. |
| BIOTEXT.dnabits.decodeFile<br>BIOTEXT.dnabits.decodefile<br>BIOTEXT.dnabits.df | Decodes a text file or a SeqRecord list with reverse DNAbits. | input_file_name: file name or list of SeqRecord;<br>output_file_name: the name for the output file. If not defined, the result will only be returned as a variable;<br>verbose: if True displays progress. | dec_list: string list;<br>if defined output_file_name a file will be saved. |
| BIOTEXT.fastatools.list2SeqRecord<br>BIOTEXT.fastatools.list2seqrecord | Converts a list of strings to a list of SeqRecord. | seq: list of biological sequences in string format; | records: list of SeqRecord. |
| BIOTEXT.fastatools.list2bioSeqRecord<br>BIOTEXT.fastatools.list2bioseqrecord<br>BIOTEXT.fastatools.list2fasta | | header: list of headers in string format, if set to 'None' the headers will be automatically defined with an increasing number. | |
| BIOTEXT.fastatools.fasta<br>Read BIOTEXT.fastatools.fastaread | Uses biopython to import a FASTA file. | input_file_name: input fasta file name. | records: list of SeqRecord. |
| BIOTEXT.fastatools.fasta<br>Write BIOTEXT.fastatools.fastawrite | Create a file using a SeqRecord list. | records: list of SeqRecord;<br>output_file_name: output fasta file name. | records: a file is saved with the defined name. |
| BIOTEXT.fastatools.getHeader<br>BIOTEXT.fastatools.getheader | Get the header from a list of SeqRecord. | records: list of SeqRecord. | headers: list with headers. |
| BIOTEXT.fastatools.getSeq<br>BIOTEXT.fastatools.getseq | Get the string from a list of SeqRecord. | records: list of SeqRecord. | seqs: list with sequences. |
| BIOTEXT.fastatools.removePattern<br>BIOTEXT.fastatools.removepattern | Removes patterns from a SeqRecord range based on regular expression. | records: list of SeqRecord;<br>rex: regular expression. | new_records: list of SeqRecord with removal applied. |

(Continue)

| | | | |
|---|---|---|---|
| BIOTEXT.fastatools.clustalOmega<br>BIOTEXT.fastatools.clustalomega<br>BIOTEXT.fastatools.clustalo | Uses the Clustal Omega to align the strings in a FASTA file. | input_file_name: input FASTA file name. | align: list with strings aligned in string format. |
| BIOTEXT.fastatools.getCons<br>BIOTEXT.fastatools.getcons | Save a temporary file with the sequences from the SeqRecord list, apply the clustalo function and obtain alignment consensus. | records: list of SeqRecord. | consensus: consensus for alignment in string format;<br>align: list with strings aligned in string format. |
| BIOTEXT.fastatools.fastatext2vect<br>BIOTEXT.fastatools.fastaText2vect<br>BIOTEXT.fastatools.fasta2vect | Perform a vectorization of a list of SeqRecord using the SWeeP, a method<br>developed by our group for FASTA vectorization. | fastatext: list of SeqRecord. | vect: matrix with the generated vectors, in ndarray** format. |
| BIOTEXT.huflus.cluster<br>BIOTEXT.huflus.clus | Perform clustering using Huflus, a method<br>developed by our group that includes the cluster number prediction. | mat: matrix in numpy.ndarray format. For example, the output of the BIOTEXT.fastatools.fastatext2vect function can be used;<br>loop: number of loops for smoothing that is used in the Huflus method, the default is 100. | cluster_idx: ndarray with the found clusters index. |

REFERENCE: Authors, 2022.

Note: *SeqRecord: Biopython object to store biological sequences and its information, as described in https://biopython.org/DIST/docs/api/Bio.SeqRecord.SeqRecord-class.html **ndarray: Numpy package object to represent array, as described in https://numpy.org/doc/stable/reference/generated/numpy.ndarray.html.

*TABLE S4. Results for the blast search using the protein WP_011156533.1*

| ID | % identity | e-value | Description | Specie |
|---|---|---|---|---|
| WP_011156533.1 | 100.000 | 1.16e-91 | hypothetical protein | Rhodopseudomonas palustris |
| WP_012494740.1 | 99.237 | 1.40e-90 | hypothetical protein | Rhodopseudomonas palustris |
| WP_107357400.1 | 96.183 | 1.24e-88 | hypothetical protein | Rhodopseudomonas palustris |
| NEW97766.1 | 96.947 | 1.66e-88 | hypothetical protein | Rhodopseudomonas sp. BR0G17 |
| NEV79437.1 | 96.947 | 3.42e-88 | hypothetical protein | Rhodopseudomonas sp. BR0C11 |
| WP_013500942.1 | 93.130 | 8.92e-86 | hypothetical protein | Rhodopseudomonas palustris |
| WP_107343178.1 | 94.656 | 1.65e-85 | hypothetical protein | Rhodopseudomonas palustris |
| WP_047308448.1 | 91.603 | 2.95e-85 | hypothetical protein | Rhodopseudomonas palustris |
| NEW90013.1 | 91.603 | 2.35e-84 | hypothetical protein | Rhodopseudomonas sp. WA056 |
| NEW92855.1 | 88.550 | 1.82e-81 | hypothetical protein | Rhodopseudomonas sp. BR0M22 |
| WP_110787918.1 | 89.313 | 5.04e-81 | hypothetical protein | Rhodopseudomonas palustris |
| WP_114358707.1 | 88.550 | 8.27e-80 | MULTISPECIES: hypothetical protein | Rhodopseudomonas |
| WP_092681992.1 | 76.923 | 3.42e-66 | hypothetical protein | Rhodopseudomonas pseudopalustris |
| WP_011439953.1 | 77.519 | 3.99e-66 | hypothetical protein | Rhodopseudomonas palustris |
| WP_011501608.1 | 76.923 | 7.97e-66 | hypothetical protein | Rhodopseudomonas palustris |
| WP_022723522.1 | 73.846 | 1.28e-64 | hypothetical protein | Rhodopseudomonas sp. B29 |
| WP_054161643.1 | 76.154 | 6.84e-62 | hypothetical protein | Rhodopseudomonas sp. AAP120 |
| WP_111360672.1 | 61.832 | 2.62e-52 | hypothetical protein | Rhodoplanes elegans |
| WP_108683060.1 | 63.780 | 2.71e-52 | hypothetical protein | Methyloceanibacter sp. wino2 |
| WP_116132402.1 | 61.069 | 3.85e-52 | hypothetical protein | Tropicimonas sp. IMCC34043 |
| WP_011474719.1 | 62.992 | 5.06e-52 | hypothetical protein | Rhodopseudomonas palustris |
| WP_011665752.1 | 60.000 | 6.30e-52 | hypothetical protein | Rhodopseudomonas palustris |
| WP_069443161.1 | 62.992 | 1.22e-51 | hypothetical protein | Methyloceanibacter stevinii |
| WP_142861444.1 | 60.305 | 1.52e-51 | hypothetical protein | Methylosinus sporium |
| WP_108916584.1 | 60.305 | 1.85e-51 | hypothetical protein | Methylosinus sporium |
| WP_110806343.1 | 61.069 | 2.46e-51 | hypothetical protein | Rhodobacter viridis |
| WP_085770093.1 | 60.630 | 1.91e-50 | hypothetical protein | Methylocystis bryophila |
| | | | | (Continue) |
| WP_132695580.1 | 59.843 | 2.70e-50 | hypothetical protein | Rhodovulum steppense |

| | | | | |
|---|---|---|---|---|
| WP_018264372.1 | 59.843 | 3.09e-50 | hypothetical protein | Methylosinus sp. LW4 |
| WP_036289079.1 | 59.843 | 3.09e-50 | hypothetical protein | Methylosinus sp. PW1 |
| WP_138166377.1 | 61.417 | 3.46e-50 | hypothetical protein | Methylocystis sp. B8 |
| KAF0130244.1 | 59.843 | 3.87e-50 | nitrogenase-associated protein | Methylocystaceae bacterium |
| WP_014889984.1 | 59.843 | 3.99e-50 | hypothetical protein | Methylocystis sp. SC2 |
| WP_123175947.1 | 59.843 | 4.22e-50 | hypothetical protein | Methylocystis hirsuta |
| WP_024878936.1 | 59.843 | 4.44e-50 | MULTISPECIES: hypothetical protein | Methylocystaceae |
| WP_124738045.1 | 60.630 | 5.14e-50 | hypothetical protein | Methylocystis rosea |
| NEP86806.1 | 62.393 | 5.37e-50 | nitrogenase-associated protein | Okeania sp. SIO2C2 |
| WP_037454643.1 | 59.843 | 7.43e-50 | hypothetical protein | Skermanella stibiiresistens |
| WP_111417306.1 | 68.702 | 8.14e-50 | hypothetical protein | Rhodoplanes roseus |
| NEP77170.1 | 61.538 | 8.50e-50 | nitrogenase-associated protein | Okeania sp. SIO3B3 |
| NEO54746.1 | 62.393 | 9.26e-50 | nitrogenase-associated protein | Okeania sp. SIO3B5 |
| WP_159730261.1 | 59.843 | 1.14e-49 | hypothetical protein | Methylosinus sp. Ce-a6 |
| JC4746 | 59.690 | 1.20e-49 | hypothetical 15k protein - rice | Oryza sativa |
| WP_036287787.1 | 57.480 | 1.38e-49 | hypothetical protein | Methylocystis sp. ATCC 49242 |
| WP_109328306.1 | 66.379 | 1.57e-49 | hypothetical protein | Azospirillum sp. CFH 70021 |
| WP_109026782.1 | 59.055 | 1.79e-49 | MULTISPECIES: hypothetical protein | Methylocystis |
| WP_017716702.1 | 60.000 | 2.06e-49 | hypothetical protein | Oscillatoria sp. PCC 10802 |
| WP_124142967.1 | 61.538 | 2.06e-49 | nitrogenase-associated protein | Okeania hirsuta |
| WP_029651706.1 | 59.055 | 2.06e-49 | hypothetical protein | Methylocystis sp. SB2 |
| WP_083240244.1 | 64.957 | 2.13e-49 | hypothetical protein | Methyloceanibacter methanicus |
| WP_018408038.1 | 59.055 | 2.13e-49 | hypothetical protein | Methylocystis rosea |
| WP_086758352.1 | 64.103 | 2.19e-49 | MULTISPECIES: nitrogenase-associated protein | unclassified Nostoc |
| WP_073598916.1 | 64.103 | 2.29e-49 | hypothetical protein | Hydrococcus rivularis |
| PPD46134.1 | 57.692 | 2.39e-49 | hypothetical protein CTY15_01795 | Methylocystis sp. |
| HAG82093.1 | 62.185 | 2.53e-49 | hypothetical protein | Cyanobacteria bacterium UBA12227 |
| NEN90905.1 | 61.538 | 2.93e-49 | nitrogenase-associated protein | Okeania sp. SIO3H1 |
| WP_096594850.1 | 64.957 | 3.32e-49 | nitrogenase-associated protein | Calothrix sp. NIES-2098 |
| WP_113892645.1 | 60.630 | 3.63e-49 | hypothetical protein | Roseiarcus fermentans |
| NET29449.1 | 61.538 | 3.89e-49 | nitrogenase-associated protein | Okeania sp. SIO1I7 |

(Continue)

| | | | | |
|---|---|---|---|---|
| NES67966.1 | 60.684 | 4.07e-49 | nitrogenase-associated protein | Okeania sp. SIO2D1 |

| | | | | |
|---|---|---|---|---|
| MTJ12972.1 | 60.684 | 5.44e-49 | nitrogenase-associated protein | Anabaena sp. UHCC 0187 |
| WP_161912950.1 | 59.055 | 6.56e-49 | hypothetical protein | Methylosinus sp. C49 |
| WP_015126499.1 | 62.393 | 6.98e-49 | nitrogenase-associated protein | Calothrix sp. PCC 7507 |
| WP_026610005.1 | 64.706 | 8.77e-49 | hypothetical protein | Methylocaldum szegediense |
| WP_003610087.1 | 57.031 | 1.12e-48 | MULTISPECIES: hypothetical protein | Methylosinus |
| NER79243.1 | 58.678 | 1.14e-48 | nitrogenase-associated protein | Leptolyngbya sp. SIO1D8 |
| WP_015210749.1 | 63.248 | 1.66e-48 | nitrogenase-associated protein | Cylindrospermum stagnale |
| NER08466.1 | 60.684 | 1.82e-48 | nitrogenase-associated protein | Okeania sp. SIO3C4 |
| WP_149468965.1 | 58.915 | 1.85e-48 | hypothetical protein | Roseomonas genomospecies 6 |
| WP_015079413.1 | 60.684 | 2.18e-48 | thioredoxin | Anabaena sp. 90 |
| WP_145622405.1 | 58.140 | 2.27e-48 | hypothetical protein | Azospirillum brasilense |
| WP_044431862.1 | 58.268 | 2.34e-48 | hypothetical protein | Skermanella aerolata |
| WP_133718238.1 | 61.240 | 2.42e-48 | hypothetical protein | Methylocaldum sp. 0917 |
| NEQ71961.1 | 58.974 | 2.82e-48 | nitrogenase-associated protein | Okeania sp. SIO2C9 |
| WP_134077353.1 | 60.630 | 3.13e-48 | hypothetical protein | Rhodovulum visakhapatnamense |
| MTJ55644.1 | 59.829 | 3.39e-48 | nitrogenase-associated protein | Anabaena sp. UHCC 0253 |
| NET45175.1 | 60.684 | 3.71e-48 | nitrogenase-associated protein | Okeania sp. SIO2B3 |
| WP_045368263.1 | 60.976 | 3.77e-48 | hypothetical protein | Methyloceanibacter caenitepidi |
| WP_156879325.1 | 59.231 | 4.14e-48 | hypothetical protein | Methylocaldum sp. BRCS4 |
| NEP04475.1 | 60.684 | 4.28e-48 | nitrogenase-associated protein | Okeania sp. SIO4D6 |
| MTJ09605.1 | 58.974 | 4.40e-48 | nitrogenase-associated protein | Anabaena sp. UHCC 0204 |
| WP_019499581.1 | 61.538 | 6.36e-48 | hypothetical protein | Pseudanabaena sp. PCC 6802 |
| WP_069968843.1 | 60.169 | 6.97e-48 | hypothetical protein | Desertifilum sp. IPPAS B-1220 |
| MTJ25652.1 | 58.974 | 7.23e-48 | nitrogenase-associated protein | Dolichospermum planctonicum UHCC 0167 |
| WP_086135375.1 | 60.465 | 8.81e-48 | hypothetical protein | Methylocaldum sp. SAD2 |
| WP_083620450.1 | 59.829 | 8.86e-48 | hypothetical protein | Planktothrix serta |
| WP_007357421.1 | 60.504 | 9.36e-48 | MULTISPECIES: hypothetical protein | Kamptonema |
| WP_011613151.1 | 59.829 | 1.17e-47 | nitrogenase-associated protein | Trichodesmium erythraeum |
| WP_017315443.1 | 61.538 | 1.19e-47 | nitrogenase-associated protein | Mastigocladopsis repens |
| WP_097070467.1 | 58.871 | 1.47e-47 | hypothetical protein | Rhodobacter maris |
| PPT11162.1 | 55.118 | 1.54e-47 | Nitrogenase-associated protein *Nif*O | Geitlerinema sp. FC II |

(Continue)

| | | | | |
|---|---|---|---|---|
| WP_028083562.1 | 58.974 | 1.57e-47 | nitrogenase-associated protein | Dolichospermum circinale |

| | | | | |
|---|---|---|---|---|
| OBQ05589.1 | 59.829 | 1.58e-47 | nitrogenase-associated protein | Anabaena sp. LE011-02 |
| WP_016917984.1 | 57.692 | 1.61e-47 | hypothetical protein | Methylocystis parvus |
| WP_112315187.1 | 60.630 | 1.63e-47 | hypothetical protein | Rhodovulum viride |
| WP_096727879.1 | 62.393 | 1.88e-47 | nitrogenase-associated protein | Nostoc carneum |
| WP_119507507.1 | 58.140 | 1.96e-47 | hypothetical protein | Azospirillum brasilense |
| WP_039727361.1 | 58.120 | 2.26e-47 | MULTISPECIES: hypothetical protein | Cyanobacteria |
| WP_075784184.1 | 59.843 | 2.27e-47 | hypothetical protein | Rhodovulum sulfidophilum |
| WP_045873077.1 | 62.393 | 2.28e-47 | MULTISPECIES: nitrogenase-associated protein | Nostocales |

Reference: Author, 2022.

TABLE S5. Demonstration of the difference between minimal and expanded encodings

| | | |
|---|---|---|
| **original text** | hypothetical 15k protein - rice | MULTISPECIES: hypothetical protein |
| **expanded encode** | HYYPYQTHYETYICYALYSYDQYDIK YSPRYQTYEYINYSYKYSRYICYE | MYVLTYISPYECYIYESYPTYSHYYPYQTH YETYICYALYSPRYQTYEYIN |
| **expanded decode** | hypothetical 15k protein - rice | multispecies: hypothetical protein |
| **minimal encode** | HYYPYQTHYETYICYALYSYDYDKY SPRYQTYEYINYSYKYSRYICYE | MYVLTYISPYECYIYESYPYSHYYPYQTHY ETYICYALYSPRYQTYEYIN |
| **minimal decode** | hypothetical 99k protein - rice | multispecies. hypothetical protein |

REFERENCE: Authors, 2022.
Note: The expanded version allows coding that maintains details of numbers and punctuation. Note change in numbers and punctuation mark in the minimal version, representing the information sensitivity loss.

*FIGURE S1. Text mining processes*

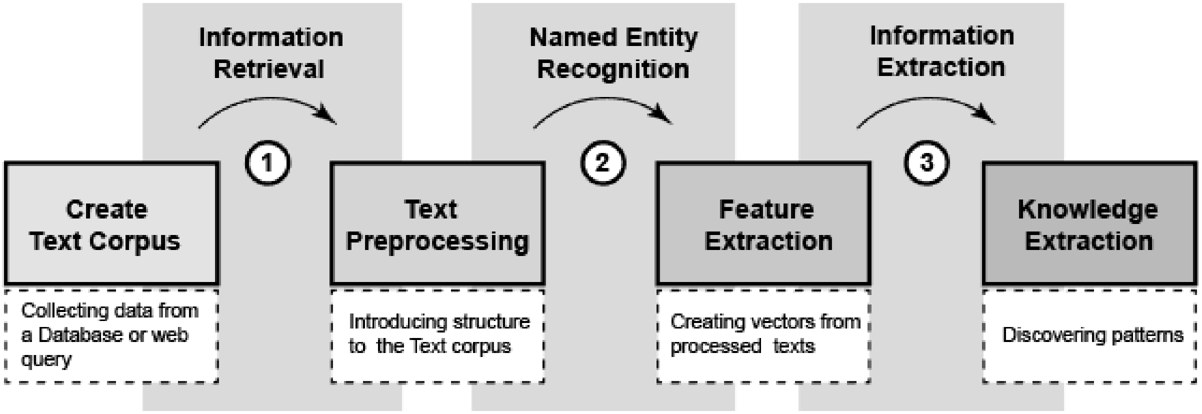


REFERENCE: Authors, 2022.
Legend: (1) Information Retrieval (IR), performed to obtain documents relevant to a particular subject of interest; (2) Named Entity Recognition (NER), which analyzes the occurrence of specific keywords in documents; (3) Information extraction (IE).

FIGURE S2. BIOTEXT strategy

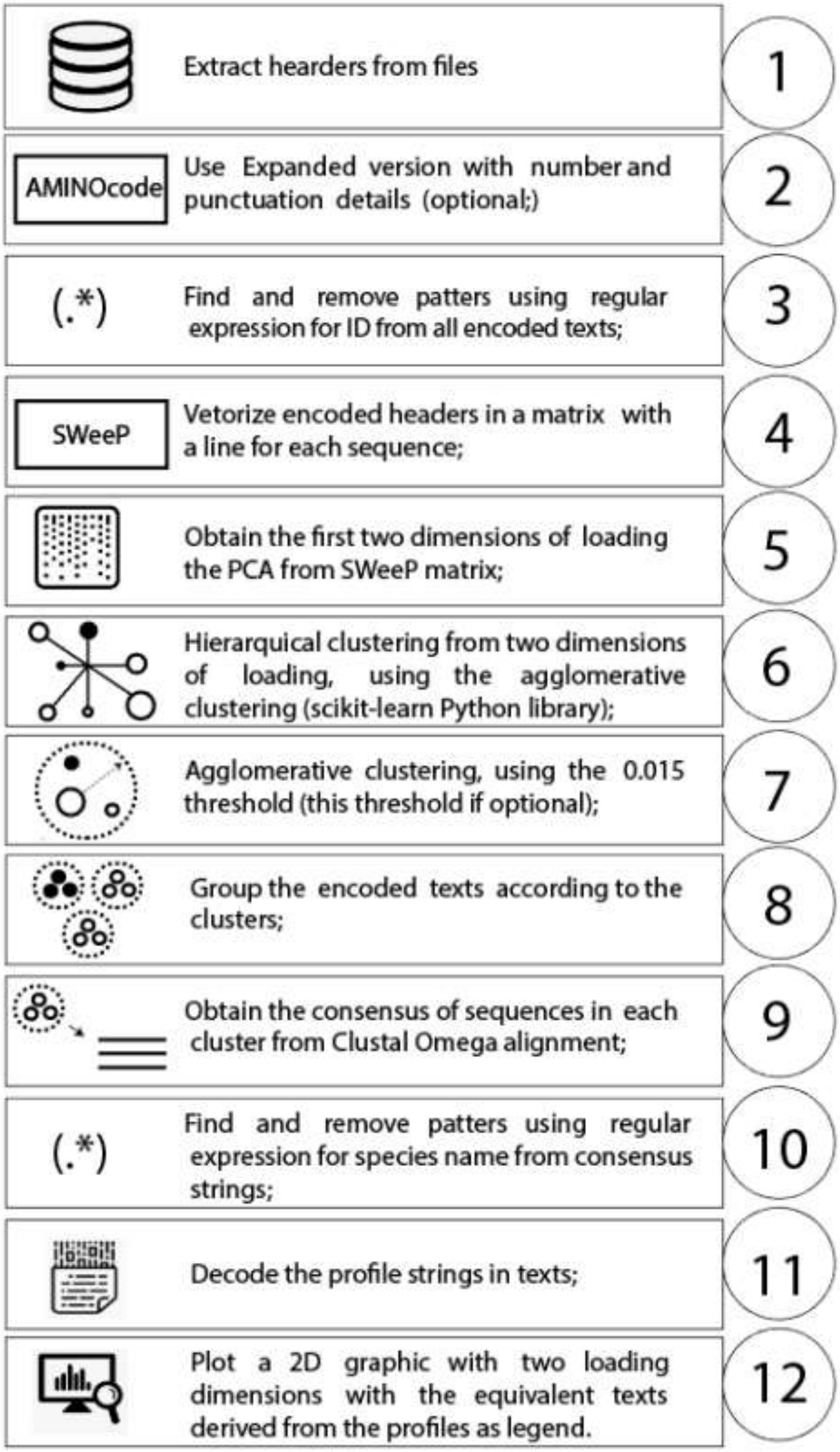


REFERENCE: Authors, 2022.

FILE S1.

Literature review details:

a) Description of the application;

b) Purpose for the field of biology, classified into 4 categories:

- General Purpose: Tools focused on describing/predicting events presented in texts in general, that can be applied for many purposes;
- Biocuration: Tools specialized in biocuration task, matching genes, proteins and associating them with functions;
- Molecular function: Tools that describe/predict molecular events, associating genes/mutations/diseases/organisms to the query texts;
- Chemicals: Tools that mainly describe/predict events related to chemicals/drugs

b) TM Tasks, according to Fleuren and Alkema (2015), which are:

- Information Retrieval (IR)
- Named Entity Recognition (NER)

Information Extraction (IE);

d) Input:

- Lists (genes, terms)
- Query (Keywords, sentences or concepts)
- Abstracts (generally, in PubMed/MEDLINE format)

Full-article and Texts (XML format);

e) Output: Each tool has its own output, depending on the type of analysis. In most cases, it consists on:

- List of genes/terms/abstracts
- Relationships between concepts/terms/genes
- Interaction/Graphical networks
- Preprocessed/tagged documents
- Statistics, predictions, matrices, scores;

f) Authors and year of publication, from 1999 to 2022.
